# Biological management, rather than chemical management, promotes the interaction between plants and their microbiome

**DOI:** 10.1101/2024.02.12.579901

**Authors:** Tianci Zhao, Xiu Jia, Xipeng Liu, Jyotsna Nepal, Eléonore Attard, Rémy Guyoneaud, Krzysztof Treder, Anna Pawłowska, Dorota Michałowska, Gabriele Berg, Franz Stocker, Tomislav Cernava, J. Theo M. Elzenga, Joana Falcão Salles

## Abstract

In the face of climate change, developing sustainable agricultural practices to reduce the use of synthetic herbicides and pesticides is crucial. However, breeding for higher yields can lead to the decoupling of plant roots and beneficial rhizosphere microbes. In this study, we aim to identify potato cultivars with functional traits facilitating efficient interactions with the rhizosphere microbiome under various agricultural treatments in the field. With the results of profiling microbial communities with amplicon sequencing data of bacteria (16S rRNA gene fragments) and fungi (ITS2 region), a piecewise structural equation model was developed. This model explains the trade-off effects of agricultural management and potato cultivars on plant growth by affecting the rhizosphere microbiome. Furthermore, we highlight that plant cultivar and the rhizosphere microbiome together determine plant below-ground growth under biological management. In contrast, both components are found to be uncoupled under chemical and control management. Our study reveals the importance of considering microbiomes in the breeding process to achieve the goals of sustainable agriculture.

## 1. Introduction

To let food production keep pace with rapid human population growth, the agricultural sector has mostly relied on technological advances that sustain plant yield. Conventional plant breeding has focused on increased productivity, maximizing yields to feed the growing global population. In several breeding programs, plant selection focuses primarily on manipulating plant traits to achieve high nutrient utilisation, rapid growth, intense stress resistance and high quality and yield (Breseghello and Coelho, 2013; Wei and Jousset, 2017). In turn, to ensure high plant production, conventional agricultural practices rely on continuous supply of chemical inputs (Timsina, 2018), which impact the environment negatively by contaminating surface water and groundwater (Ju et al., 2006; Goulson, 2013), increase greenhouse gas emissions (Snyder et al., 2009; Gao and Cabrera Serrenho, 2023), reduce soil microbial biomass and activity (AL-Ani et al., 2019), and already changed the plant microbiome (Berg and Cernava, 2022). Under the current climate change challenges, minimising the negative impacts of agriculture to achieve sustainable and environment-friendly crop production is essential. However, sustainable agriculture depends not only on the willingness of farmers to reduce chemical inputs but also on the availability of plant genotypes that can efficiently use the already existing soil nutrients for growth.

As the fundamental substrate for plant growth, the soil provides water and essential nutrients and plays a vital role in supporting plant growth through the activity of soil microbes. Numerous studies have confirmed that soil microbiota have beneficial effects on plant development. Soil microorganisms can facilitate plant growth by promoting nutrient uptake (Zaidi and Khan, 2005; van der Heijden et al., 2008) or by regulating phytohormone metabolism, thus regulating plant growth, stimulating root branching and plant defence (Krome et al., 2010; Van Der Meij et al., 2023). Root-associated microorganisms also play a role in attenuating the plant response to biotic stress by increasing plant resistance (Santhanam et al., 2015; Chauhan et al., 2019) or by inducing systemic resistance through the production of bacterial volatile organic compounds (Ryu et al., 2004; Garbeva and Weisskopf, 2020). Root-associated microorganisms can contribute to alleviating the effects of abiotic stress on plants, for instance by improving water uptake and nutrient accumulation (Köhl et al., 2016; Mbodj et al., 2018) or by increasing the level of protective compounds such as ascorbate and proline, during drought stress (Ruíz-Sánchez et al., 2011). Given the potential of soil and plant microbiomes for supporting plant development, harnessing mutualistic interactions between microbiomes and plants should become a priority in our breeding programs. However, the beneficial functions of soil microbes on plant development are hardly being considered in plant breeding (Wei and Jousset, 2017; Wille et al., 2019). For such an approach, selecting plant traits that stimulate the interaction with beneficial soil microbes is necessary (Kusstatscher et al., 2021).

Plant functional traits have been shown to impact the composition of microbial communities (Legay et al., 2016). It is well-established that plant root exudates represent an important nutrient source for soil microorganisms that colonize the rhizosphere (Reinhold-Hurek et al., 2015) and that the secondary metabolites produced by plant roots can alter the composition of the rhizosphere microbial community and enhance plant defence mechanisms (Hu et al., 2018). Although less studied, root traits have been demonstrated to correlate with the fungi-to-bacteria (F/B) ratio in the soil and plays a role in carbon and nitrogen cycling (Legay et al., 2014; Wan et al., 2021). Given the importance of the chemical and physical components of the roots, we have recently selected a subset of seven potato cultivars showing potential for diverse levels of microbiome interactive traits (MITs) such as morphological root traits, root exudation (quality and quantity of exudates), and microbial diversity in the rhizosphere under control conditions (Zhao et al greenhouse paper). Nevertheless, there is limited information regarding the performance of these cultivars with MITs under real-world conditions.

To address this question, a field trial was conducted aimed at determining how potato cultivars with different MITs recruit and interact with soil microbes in field conditions under different agricultural management regimes, which impact soil physical and chemical parameters, thereby affecting the interactions between plants and microbes (Schmidt et al., 2019). Various treatments were implemented in the field trial: chemical treatments involved the application of synthetic fertilizers and pesticides, while biological treatments entailed the introduction of microbial strains. Hence, we explored the ability of the selected seven potato cultivars (including a commercial one, Desiree) to interact with the soil microbiome under different treatments. We hypothesize that the selected potato cultivars with high MITs can shape or interact with healthy and beneficial microbiomes in the rhizosphere and are, hence, less dependent on chemical inputs. We further hypothesize that agricultural treatments will shape the rhizosphere microbial community, thereby affecting plant performance. Moreover, we expect that biological treatments will further stimulate the rhizosphere microbiome and plant growth, an effect that will be more prominent in MIT-selected cultivars.

## 2. Materials and methods

### 2.1. Cultivar selection

The potato (*Solanum tuberosum* L.) cultivars included in this study were carefully selected through a stepwise process based on MITs associated with root traits, quality and quantity of root exudation, and bacterial and fungal rhizosphere diversity (Zhao et al greenhouse paper). Out of thousands of cultivars, seven cultivars with a diverse performance of the above characteristics are chosen to be tested under field conditions: Atol, Desiree, Jelly, Krab, Pasja Pomorska, Rudawa and Salto. For comparison purposes, we use the commercial cultivar Desiree as a reference.

### 2.2. Field trial setup

From June to October 2021, we performed the experiment in a field near Valthermond, the Netherlands (52°51’8.3”N, 6°55’5.2”E). The field had been used for potato cultivation in 2017 and 2018 and maize cultivation in 2019 and 2020, after using grassland as dairy farm for about 20 years. In the field trial, six treatments were performed (Fig. S1). Specifically, the **Control** treatment is a minimum pesticide input to maintain potato growth from Colorado potato beetles (*Leptinostarsa decemlineata*). The **Consortium B** treatment consists of *Bacillus* sp., *Pseudomonas* sp., and *Stenotrophomonas* sp. in a 10^10^ CFU/ml concentration per isolate. The bacterial consortium was added in two steps. In the first step, we dipped the tubers in the consortium (diluted to a concentration of 10^8^ CFU/ml) on planting day. In the second step, we added 10^9^ CFU per plant of the bacterial consortium when the shoots of most plants were visible on the 22^nd^ day after planting. The **Consortium BP** treatment contained a mixture of plant-growth promoting bacteria (ExSol P: 10^9^ CFU/ml of *Bacillus amyloliquefaciens* ECO B 01 (DSM 32047) and *Bacillus pumilus* ECO B 02 (DSM 32048), protozoa (ECO – P01 – P02: 10^5^ cysts/ml of *Cercomonas lenta* ECO P 01 (DSM 32401) and *Rosculus terrestris* ECO P 02 (DSM 32402). Each potato plant received 10^7^ CFU *Bacillus* sp. and 10^4^ cysts protozoa on the 22^nd^ day after planting. The **Fertiliser** treatment was performed with 35 kg/ha potassium phosphate addition during the planting day. The **Pesticide** treatment was set against potato late blight driven by *Phytophthora infestans*. The fungicide propamocarb (625 g/l) and herbicide prosulfocarb (800 g/l) were the main compounds (Table S1). The **Fertiliser plus Pesticide** treatment combined the previous two treatments. It represents a conventional method farmers use when growing potatoes. In each treatment, three replicate plots were set up. The treatments with Consortium B and Consortium BP are referred to as **Biological** management. The Fertiliser, Pesticide, and Fertiliser plus Pesticide treatments are referred to as **Chemical** management. The Control treatment is referred to as **Control** management.

### 2.3. Plant performance

The plants were harvested at the fifth week after planting. Plant performance was assessed via measurements of plant height, plant above- and below-ground biomass. Root-to-shoot biomass ratio was used as an indicator of plant root traits. Plant above- and below-ground dry weight were determined after drying corresponding plant materials at 60 °C for 48 hours. We collected three subsamples (from three plant individuals) in each replicate plot and combined them into one pooled sample. In this case, even though only three replicates were set, nine sub-replicates can be found in the figures of plant data.

### 2.4. Soil sampling and sequencing

Before planting, six pooled bulk soil samples (day 0) from six treatment blocks were collected. In the fifth week after planting, the same as plant data, we collected 126 pooled rhizosphere soil samples (7 cultivars × 6 treatments × 3 replicate plots). The rhizosphere soil was washed from plant roots with a 1 × phosphate buffer and Silwet® L-77, following the protocol from Lundberg et al. (2012). Briefly, the tube containing roots and buffer was vortexed, and the solution was filtered through a 100 µm nylon mesh cell strainer. The turbid filtrate was then centrifuged to collect the sediment as rhizosphere soil. We also collected 18 bulk soil samples (6 treatments × 3 replicate plots) in the fifth week. Overall, 150 soil samples were collected for processing.

DNA from bulk and rhizosphere soil was extracted using the DNeasy PowerSoil kit (Qiagen, Hilden, Germany) on 0.25 g of soil. DNA extraction followed the kit’s instructions, except for the first step of bead beating with FastPrep-24™ 5G Instrument at 6000 rpm/s for 40 s (MP Biomedicals, Santa Ana, USA). The bacterial community was profiled by targeting the V4 region of the 16S rRNA gene using the forward primer 515F (5′-GTGCCAGCMGCCGCGGTAA-3′) and reverse primer 806R (5′-GGACTACHVGGGTWTCTAAT-3′) (Caporaso et al., 2011, 2012). The ITS2 region was sequenced for the soil fungal community using the forward primer 5.8SR (5’-TCGATGAAGAACGCAGCG-3’) and reverse primer ITS4 (5’-TCCTCCGCTTATTGATATGC-3’) (White et al., 1990). The Illumina MiSeq platform was used for paired-end sequencing (2 × 250bp for 16S, 2 × 300bp for ITS) at the PGTB (Genome Transcriptome Platform of Bordeaux, doi:10.15454/1.5572396583599417E12) (Univ. Bordeaux, INRAE, BIOGECO, F-33610 Cestas, France).

### 2.5. Sequencing data processing

16S rRNA gene sequencing data were analysed with the next-generation microbiome bioinformatics platform QIIME 2 / 2020.8 (Bolyen et al., 2019). After removing adaptors and primer sequence with Cutadapt (Martin, 2011), the sample sequence was filtered, denoised, and dereplicated using Divisive Amplicon Denoising Algorithm (DADA2) with default setting (Callahan et al., 2016). The 16S rRNA sample reads’ taxonomic classification was obtained through the SILVA database (version 138) (Yilmaz et al., 2014). The PIPITS / 2.8 pipeline performed ITS sequencing data (Gweon et al., 2015). Taxonomic assignment was approached with the UNITE database (version 8.3) (Kõljalg et al., 2013).

### 2.6. Statistical Analysis

The two-way analysis of variance (ANOVA) was used for testing the variations in plant performance. Duncan’s or Tukey’s honest significant differences multiple range tests were applied as post hoc comparisons. The soil microbial community analysis (bacteria and fungi) was performed in R (version 4.2.0). The feature tables were rarefied at 5700 reads for bacterial 16S rRNA gene sequences and 9800 for fungal ITS sequences. Finally, we identified 5422 amplicon sequence variants (ASVs) for the bacterial community and 4162 operational taxonomic units (OTUs) for the fungal community. The soil microbial community structures were visualised through Principal coordinate analysis (PCoA) based on Bray-Curtis and weighted UniFrac distances. Permutational multivariate analysis of variance (PERMANOVA) was used to evaluate whether the cultivars and treatments significantly influenced the soil microbial community composition. Spearman Correlation Analysis was used to assess the correlation between microbial components.

To compare the agricultural treatment effects on the rhizosphere microbial inter-kingdom interactions. We provided the co-occurrence patterns to display the interactions between bacteria and fungi under different management practices by R package “igraph” (Csardi and Nepusz, 2006). In particular, to facilitate a comprehensive comparison of the microbial community responses to different agricultural treatments, we highlighted the treatment-sensitive ASVs/OTUs according to the workflow in cropping research (Hartman et al., 2018a). The intersection of function multipatt in package “indicspecies” (De Cáceres et al., 2010) and function glmLRT in package “edge R” (Robinson et al., 2010) defined the sensitive ASVs/OTUs. Sensitive means the ASVs/OTUs differ significantly (p < 0.05) between treatments.

To explore the interactions between plant and rhizosphere microbiomes, piecewise structural equation modelling (piecewise SEM) was used to evaluate the direct and indirect correlations among different factors. The model was constructed with the package “piecewiseSEM” (Lefcheck, 2016) and visualised with the package “semPlot”. The first axis of principal coordinate analysis (PCoA1) of microbial community beta-diversity (bacteria and fungi) was composited as one predictor to represent microbial community composition in the model (Delgado-Baquerizo et al., 2016; Tian et al., 2021). We used Shannon diversity to represent microbial community alpha-diversity. Standardized coefficients were extracted to assess the relative importance of each microbial group. Subsequently, a composite predictor, termed microbial community composition/diversity, was derived by combining bacterial PCoA1/Shannon and fungal PCoA1/Shannon using the obtained coefficients. Categorical variables (i.e., potato cultivars, treatments and management) were encoded numerically for model compatibility. Akaike’s information criterion (AIC) and the Bayesian information criterion (BIC) are used for model selection (Akaike, 1974; Schwarz, 1978). Smaller AIC or BIC suggest a better-fitting model. Fisher’s C statistic was used to test the model’s goodness of fit of the model. The model fits well when Fisher’s C is low and the *P*-value is higher than 0.05.

## 3. Results

Plant height and above- and below-ground biomass were used as proxies for the plant performance of seven cultivars. The rhizosphere soil microbial communities from six different agricultural treatments were profiled using amplicon sequencing data. Our findings revealed that plant performance is a cultivar-dependent trait, with agricultural treatments exerting limited influence. Notably, the performance of most of the plant cultivars selected for microbial interactive traits (MITs), exhibited better than that of commercial cultivar Desiree. The rhizosphere microbial community responded more strongly to agricultural treatments than potato cultivars. Under biological management (i.e., Consortium B and Consortium BP treatments), the rhizosphere microbiome strongly correlated with plant growth.

### 3.1. Plant Performance

Plant performance of the seven potato cultivars in the field experiment is significantly influenced by both cultivar and treatment (ANOVA, Table 1). Cultivar had a more substantial effect on plant growth (*F_plant height_* = 29.94, *F_above-ground biomass_* = 19.39, and *F_below-ground biomass_* = 32.38; Table 1) than treatments (*F_plant height_* = 6.547, *F_above-ground biomass_* = 6.865, and *F_below-ground biomass_* = 3.443; Table 1). While the interaction between cultivar and treatment only significantly influenced plant below-ground biomass (*p* < 0.05; Table 1). Treatment and cultivar both had a significant effect on the root-to-shoot dry weight ratio (*p* < 0.001; Table 1).

**Table 1.**
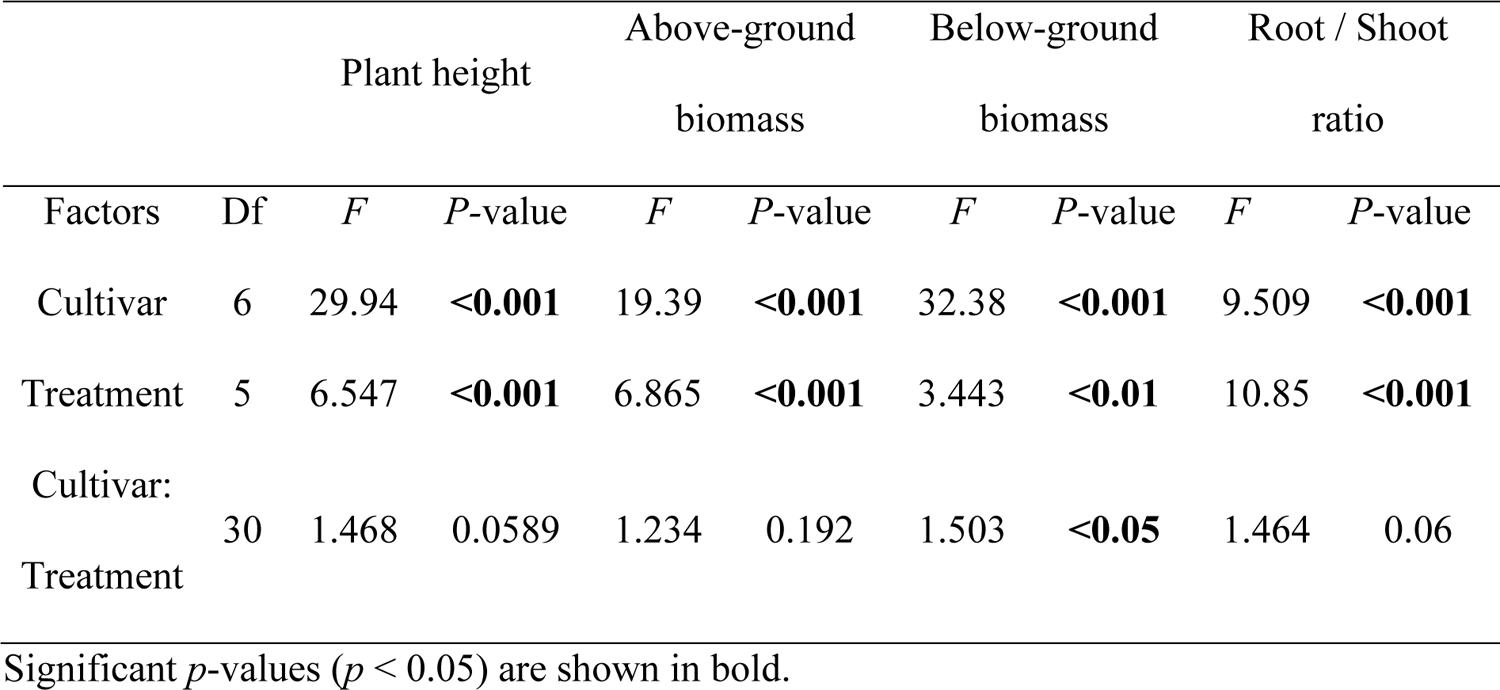
Two-way variance analysis (ANOVA) shows the influence of cultivar and treatment on plant performance: plant hight, above- and below-ground biomass, and root-to-shoot ratio.

Cultivar significantly influenced plant performance when subjected to the same treatment (Fig. 1). Regarding plant above- and below-ground growth, the cultivar Salto showed relatively high plant development in all treatments. Rudawa also performed well in the majority of treatments, except for Consortium B treatment. Pasja Pomorska performed well in Control and Pesticide treatments but showed comparatively poor performance in the Consortium B treatment. Desiree and Jelly showed less vigorous growth compared to other cultivars across all treatments. In terms of root traits, the addition of pesticides removed the root-to-shoot ratio variation among different cultivars. Salto exhibited strong root development across all treatments. Additionally, Jelly displayed a higher root-to-shoot ratio under all high-nutrient conditions (i.e., Fertiliser and Fertiliser plus Pesticide treatments).

**Fig. 1.**
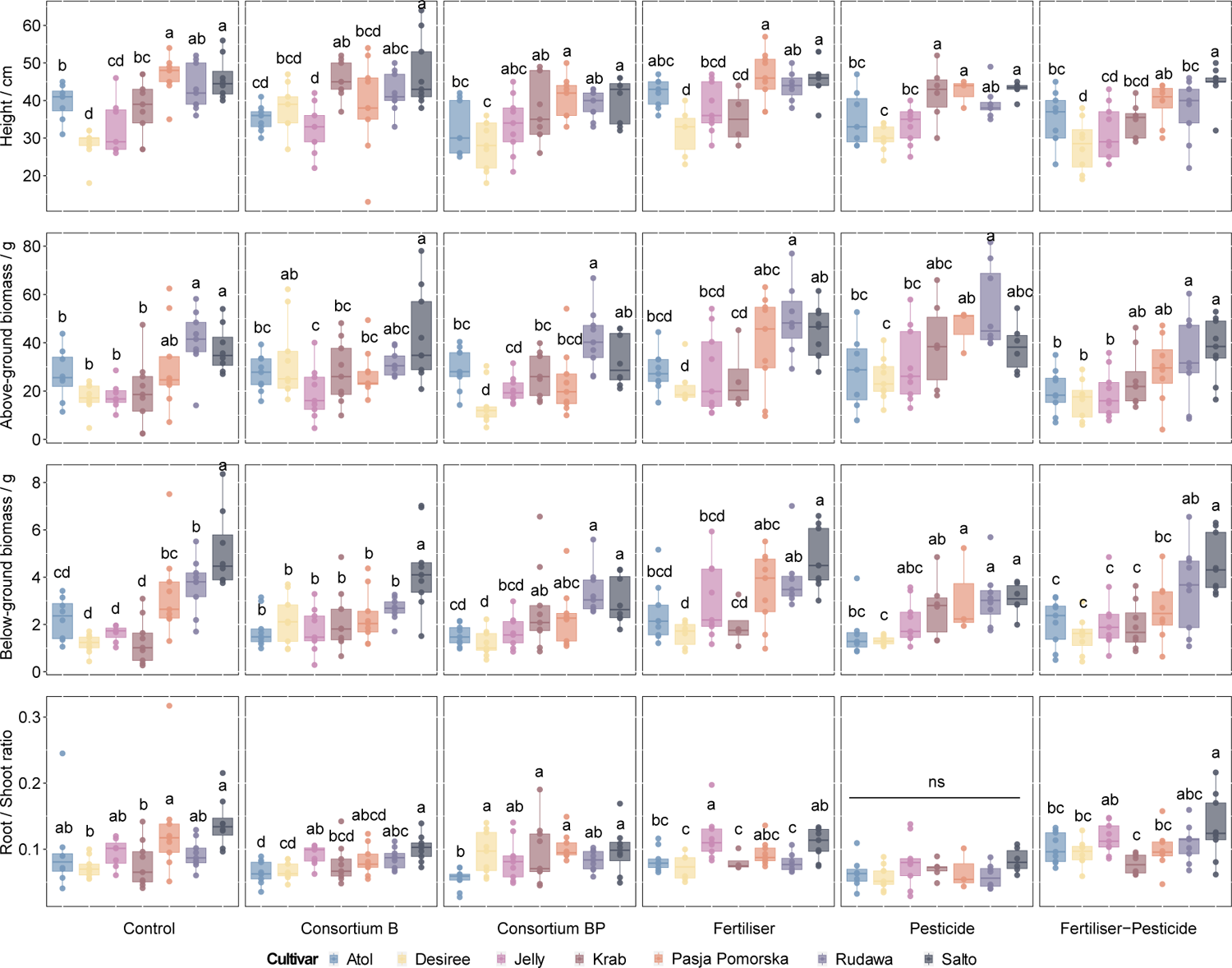
Plant performance of seven potato cultivars across treatments. Colours represent different cultivars. Root/Shoot ratio means the root-to-shoot dry weight ratio. Letters indicate significant differences across cultivars (Duncan post hoc test).

When evaluating the plant cultivar growth across different treatments (Fig. S2), Salto performed the same above-ground biomass regardless of treatments. For other plant development factors, Salto exhibited well when subjected to Control and Fertiliser plus Pesticide treatments. Treatments significantly influenced Desiree growth. In particular, Consortium B increased Desiree plant above- and below-ground growth more than other treatments.

### 3.2. Soil and Rhizosphere Microbial Community Composition

The alpha diversity of the rhizosphere soil microbial community was visualized for the different treatments (Figs. S3 and S4). There was no significant difference between different cultivars under the same treatments. The taxonomy of different kingdoms was shown by management types (Figs. S5 and S6). Slight variations were found between different cultivars and managements.

The microbial community composition of bulk soil not in direct contact with plant roots, was compared with the rhizosphere soil. Soil microbial community structures significantly differed between soil compartments (*p_bacteria_* = 0.001, *p_fungi_* = 0.001; Fig. S7). Specifically, the bacterial and fungal communities of bulk soil samples clustered together and were separate from the rhizosphere microbial community. In addition, the fungal community had the most substantial variation in soil compartments compared to the bacterial community (*R*^2^*_bacteria_* = 0.08, *R*^2^*_fungi_* = 0.17; Fig. S7).

The treatment had a much more significant impact on the composition of the rhizosphere microbial communities compared to the potato cultivars, as indicated by the PERMANOVA results (Fig. 2). The first two axes of PCoA plots also showed the separation of the rhizosphere bacterial and fungal communities by treatments, suggesting that the majority of the variation in microbial beta diversity was attributed to differences in treatments (Fig. 2a,b). In addition, the different treatments had varied effects on the microbial communities, with the fungal community responding more strongly to treatments than the bacterial community (*R*^2^*_bacteria_* = 0.16, *R*^2^*_fungi_* = 0.25; Fig. 2a,b). Specifically, the bacterial community at the Fertiliser and Fertiliser plus Pesticide treatments were clustered separate from Control compared to other treatments (Figs. 2a and S9a). The fungal community was mainly affected by the Pesticide and Fertiliser plus Pesticide treatments, which were clustered separate from Control in the fungal community (Figs. 2b and S9b). By grouping different treatments into biological management (i.e., Consortium B and Consortium BP treatments) and into chemical management (i.e., Fertiliser, Pesticide, and Fertiliser plus Pesticide treatments), we found that the microbial communities of biological management were closer to Control than chemical management (Fig. 3). The results suggested that biological management caused less disturbance on the rhizosphere microbial community structure than chemical management.

**Fig. 2.**
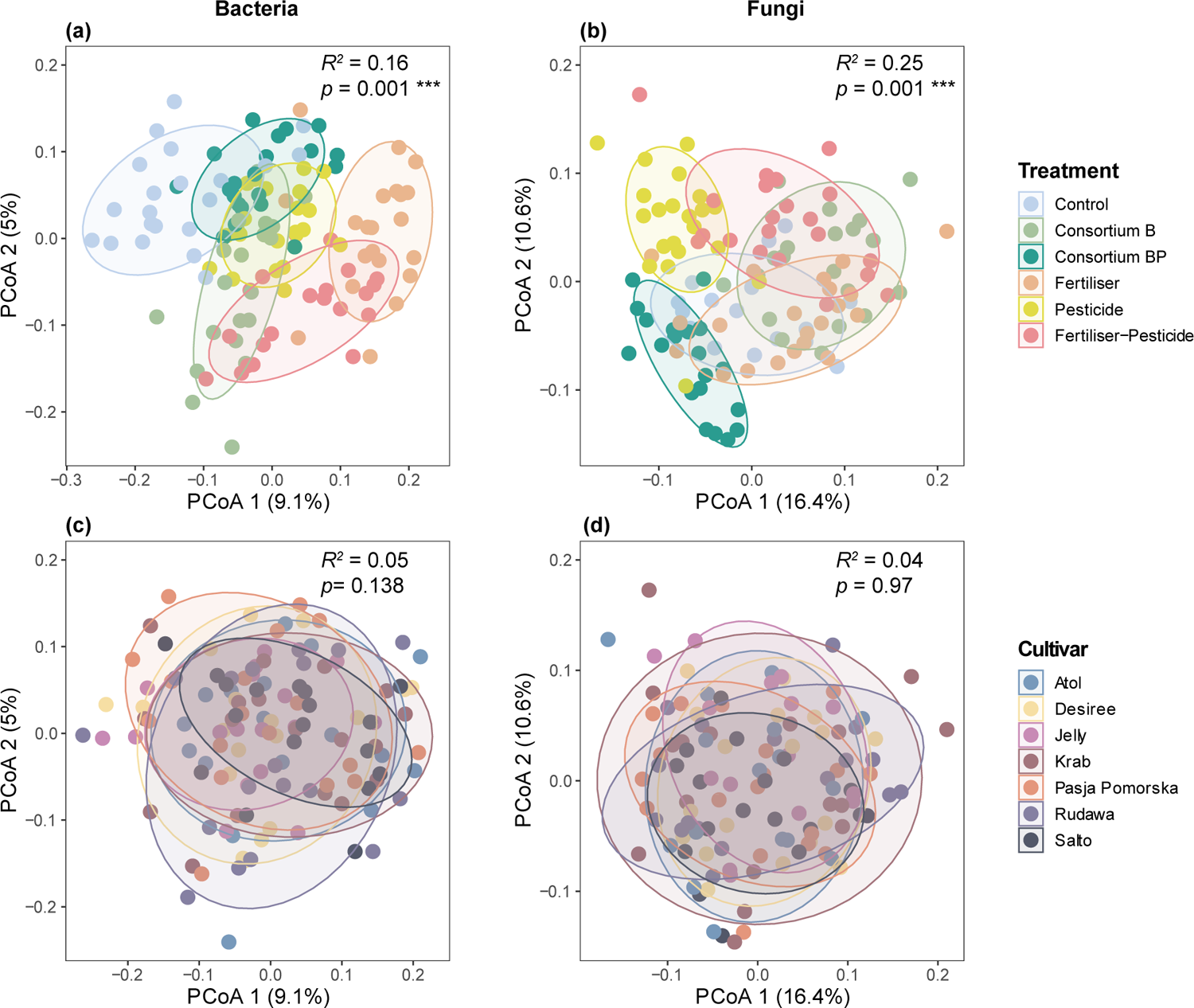
The effect of treatment and cultivar on the composition of microbial communities in rhizosphere soil. Microbial composition was visualized via a principal coordinates analysis (PCoA) based on Bray-Curtis dissimilarity matrices. The community dissimilarities of bacteria and fungi are displayed separately on the left and right, respectively. The upper panel depicts groups by treatments, while the lower panel present groups by cultivars. PERMANOVA (Adonis) results in the upper right corner of each panel show the influence of factors on community composition. *R*^2^ explains variation, and *p* values are based on 9999 permutations. *** indicates the significant *p* values (*p* = 0.001).

**Fig. 3.**
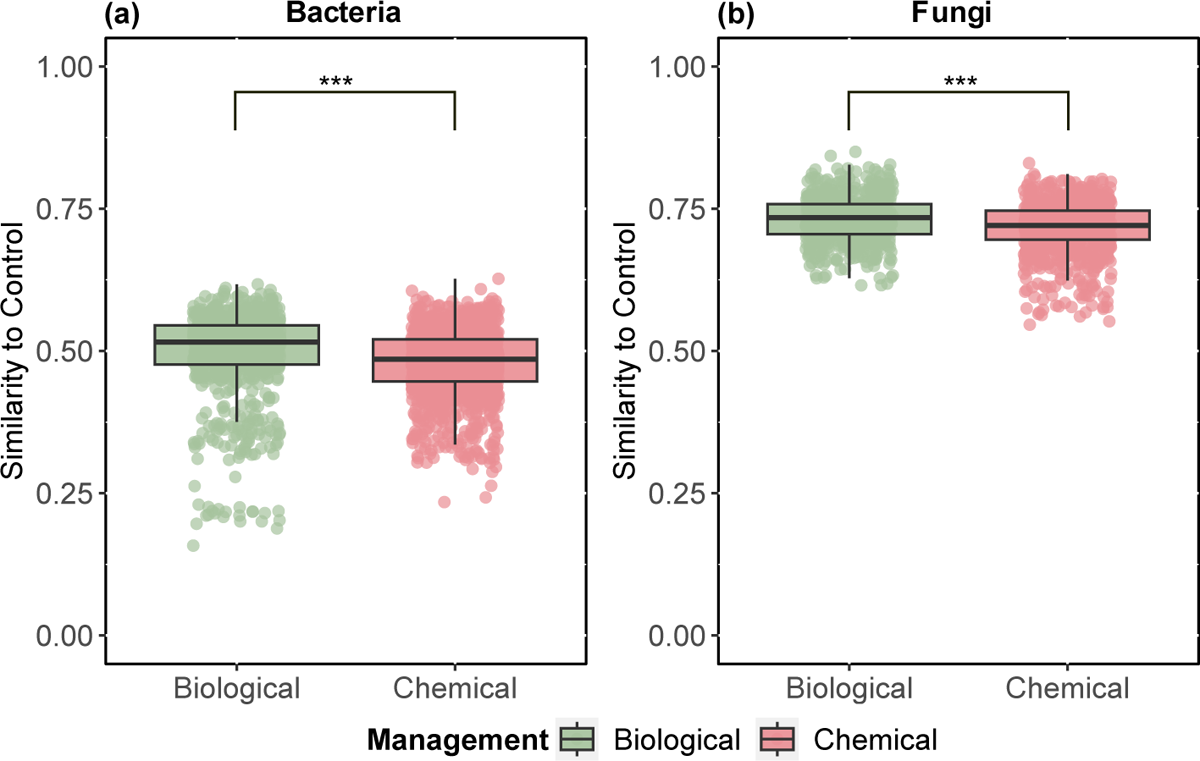
The similarity of management on the distance of rhizosphere soil microbial communities, compared to Control. Displayed for bacteria and fungi were visualised separately based on similarity, which is 1-Bray-Curtis matrices. *** indicates significant differences across management regimes (*p*<0.001; one-way ANOVA).

Regarding the potato cultivars, they didn’t have a significant impact on the structure of the rhizosphere microbial communities (Fig. 2c,d). Specifically, it was observed that Jelly displayed the lowest variation in the bacterial community across treatments, while Salto exhibited the lowest variation in the fungal community across treatments, suggesting that treatments had less effect on the rhizosphere communities of Jelly and Salto than of other potato cultivars (Fig. S10).

We further looked into the variation of microbial community composition across cultivars in each treatment. The results indicated that, except for the Fertiliser plus Pesticide treatment, the influence of cultivar on bacterial community composition was not statistically significant across all other treatments (Fig. S11). Cultivar did not significantly affect fungal community composition for all treatments (Fig. S12).

### 3.3 Agricultural management influences microbial co-occurrence patterns

We compared the co-occurrence network attributes of different management practices (Table 2). The average value for each management was calculated to make a comparison between the three different management practices possible. The result showed that biological and chemical management decreased the complexity of network connections between soil microbial communities compared to control. Specifically, the correlation links decreased from 12157 for control to 7779 for biological and 6660 for chemical management. Chemical management had the most network nodes (N_control_ = 1302, N_biological_ = 1484, N_chemical_ = 1534; Table 2) but the least network edges, suggesting that chemical management reduces the connectivity of soil microbial communities.

**Table 2.**
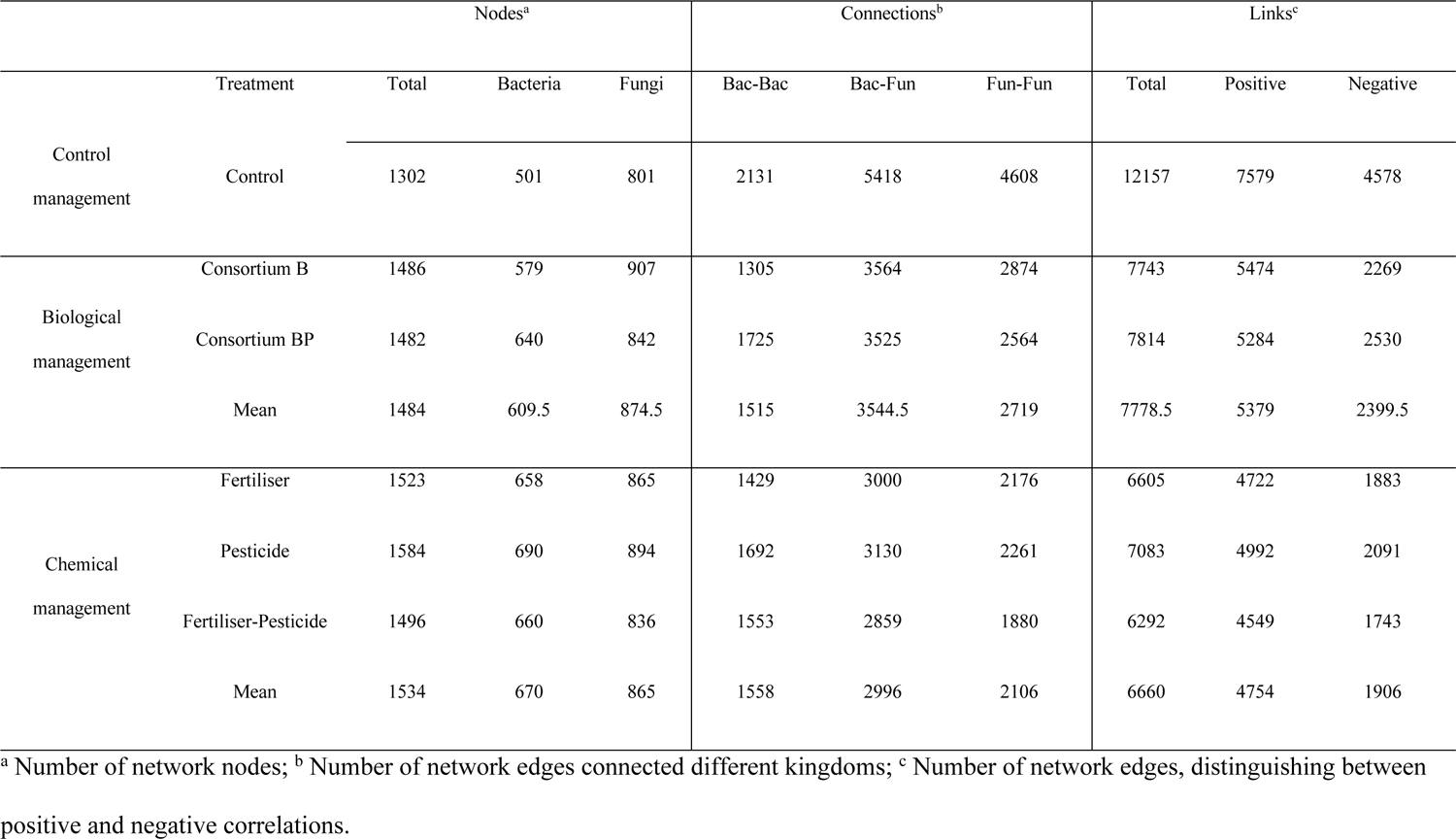
Correlation attributes of rhizosphere co-occurrence networks.

Biological management had a more intensive microbial connection than chemical management (Fig. 4). The sensitive ASVs/OTUs enriched in Consortium B treatment were clustered. In chemical management, Fertiliser treatment contained more amount of sensitive ASVs/OTUs than other chemical treatments. The number of correlation links in biological management was four times that of chemical management (1763 for biological and 483 for chemical, respectively). Chemical management had more positive connections than biological management (93% for chemical and 82% for biological, respectively). Compared to biological management, chemical management increased the proportion of bacteria-to-bacteria links and decreased the proportion of fungi-to-fungi and fungi-to-bacteria links (Fig. 4). To conclude, biological management increased the interactions between the rhizosphere microbes compared to chemical management.

**Fig. 4.**
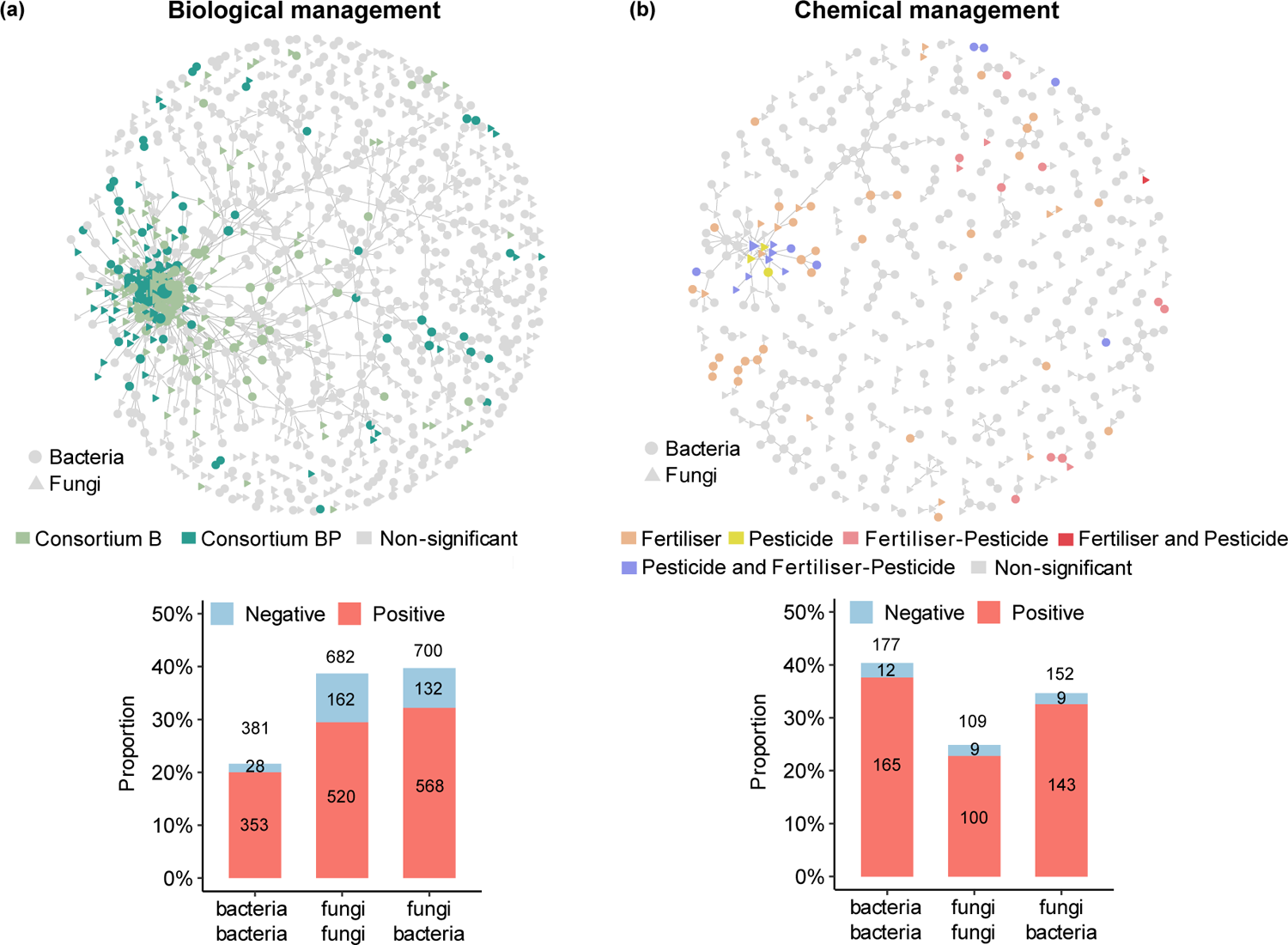
(a) and (b) Co-occurrence networks of treatment-sensitive ASVs/OTUs, displayed for biological and chemical management methods. The upper panels visualise the significant correlation (coefficient > 0.6, *p* < 0.01) between bacterial ASVs and fungal OTUs in rhizosphere soil communities. Circles and triangles refer to bacteria and fungi, respectively. ASVs/OTUs are coloured based on their association with the different treatments, indicating a significantly higher abundance in the treatment. Grey ASVs/OTUs are insensitive to treatments in management. The size of nodes indicates the degree of edges connected to the ASVs/OTUs. The lower panels show the proportion of positive and negative correlations in biological and chemical management. The numbers on each bar, from bottom to top, represent the count of positive, negative, and total links, respectively.

### 3.4. Piecewise structural equation modelling

A piecewise structural equation modelling (piecewise SEM) constructed to investigate the interactions between plant growth, cultivar and rhizosphere microbiomes under various management regimes (Fig. 5). The results revealed a direct influence of cultivar on plant below-ground growth, while management practices were shown to impact the rhizosphere microbial community. In all management practices, cultivar was the main factor affecting plant below-ground growth. Management had a significant effect on microbial composition, which was also associated with plant growth (Fig. 5a). Specifically, for control management, cultivar and microbial community diversity affected plant growth, but no significant influence was found on microbial community composition (Fig. 5b). For biological management, different treatments can affect the composition of the microbial community. The directly standardised effect level of treatment on the microbial community was negative, which means that the microbial community in the treatments were different from each other. Notably, the regulation of plant below-ground biomass was attributed to microbial community composition and plant cultivar (Fig. 5c). However, regarding chemical management, the microbial community composition was significantly impacted by treatments but unrelated to plant growth (Fig. 5d). Chemical management, when compared to biological and control management, eliminated the effect of rhizosphere microbiome on plant development, thereby decoupling the rhizosphere microbiomes from plant growth.

**Fig. 5.**
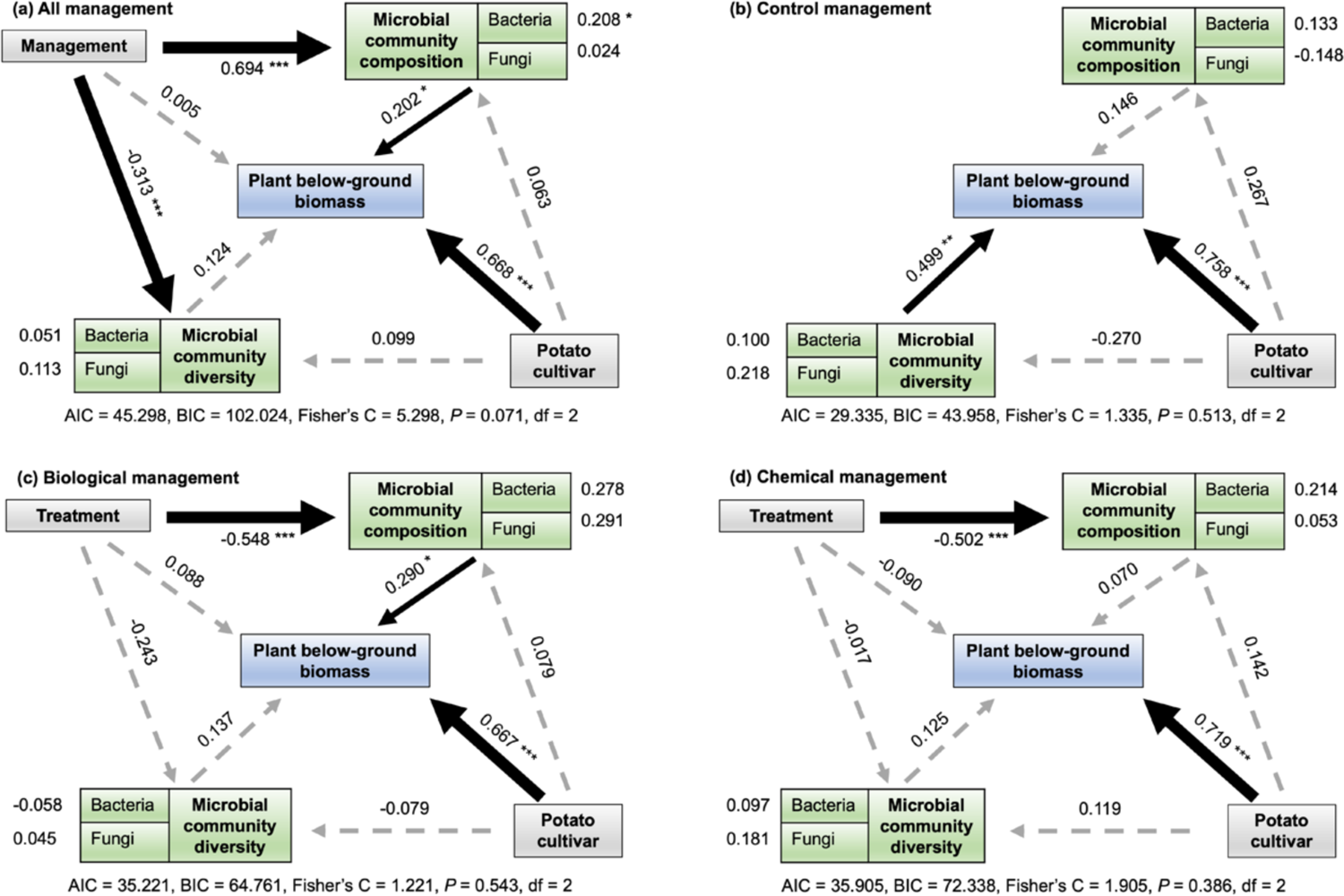
Piecewise structural equation modelling (piecewise SEM) showed direct and indirect links between rhizosphere community and plant cultivar effects on the plant below-ground biomass based on different management. Biological management contains Consortium B and Consortium BP treatments. Chemical management contains Fertiliser, Pesticide and Fertiliser plus Pesticide treatments. The dashed arrows represent non-significant correlations. The solid arrows represent significant correlations. Numbers on the arrows are path coefficients that represent the directly standardised effect level of the correlation. The thickness of the arrow represents the strength of the correlation. *, **, and *** indicate *p* < 0.05, *p* < 0.01, *p* < 0.001, respectively.

## 4. Discussion

Sustainable agricultural practices aim to eliminate the environmental footprint of agriculture by reducing the use of chemical inputs and stimulating the use of biological practices. The success of this transition relies partially on using crops capable of harnessing beneficial functions of soil microbiomes. However, current breeding processes do not select plant cultivars for optimal mutualistic interactions with soil microbes. Here we evaluate the potential to reverse the current mismatch between sustainable production and conventional cultivars by testing a selection of potato cultivars that have been screened for high microbiome interactions. By investigating plant performance and the root-associated microbiome of seven potato cultivars under different agricultural management practices, our study reached two main conclusions. First, we demonstrated that most of the plant cultivars with microbiome interactive traits (MITs) have a better plant performance than the commercial cultivar Desiree in the field. Second, we revealed that agricultural treatments strongly regulated the rhizosphere microbiome. Specifically, we showed that biological management promotes the interaction within rhizosphere microbiome and its effect on plant performance, whereas this effect is absent under chemical treatments. Below we discussed these results by placing them in the context of future breeding strategies for sustainable agricultural production.

### 4.1. MIT selection supports plant performance

Overall, we could demonstrate that the cultivars with MITs improve the capacity of plant performance compared to the commercial cultivar Desiree (Fig. 1). Specifically, Salto has notable performance in terms of plant growth even in the absence of chemical inputs. In relation to the pilot experiment, Salto exhibits a high root-to-shoot biomass ratio and root exudate metabolite level (Zhao et al greenhouse paper). According to the European cultivated potato database (https://www.europotato.org/), Salto is characterised by its high resistance to pests, in the context with our results, indicating that the plant traits of Salto are adaptable to various conditions and have the potential to interact better with soil microbiome to facilitate plant growth. Our results corroborate the importance of selecting plant cultivars with favourable root traits as breeding targets to bolster plant resilience in the face of climate change (Ober et al., 2021). This study reveals the importance of MITs in promoting plant-microbiome interactions, and their influence on plant growth, ultimately providing a new strategy for breeding.

Although the approach used in this study showed the potential of identifying potato cultivars with higher MIT, further investigation is needed to identify those traits at the plant genome level and use them in breeding programs. Genome-wide association studies (GWAS) have been widely used to link plant traits to specific traits (Tibbs Cortes et al., 2021). Several quantitative trait loci (QTL) were shown to shape the root traits of durum or common wheat to influence the host agronomic performance (Canè et al., 2014; Voss-Fels et al., 2018; Li et al., 2020). A study used leaf metabolite (GC-MS) and transcript markers (qRT-PCR) datasets of 31 potato cultivars during early developmental stages to select drought tolerance gene markers to predict field yield under drought stress, thus reducing the cost of breeding trials (Sprenger et al., 2018). In line with our results, utilising gene expression and genome sequencing to identify QTLs associated with MIT that can effectively interact with the soil microbiome can provide a feasible breeding strategy. This approach can help pinpoint specific genes or genetic regions responsible for beneficial plant-microbiome interactions, contributing to the development of crop cultivars with enhanced performance and adaptability.

### 4.2. Agricultural management drives the composition of the rhizosphere microbiome

Although cultivars are important for plant growth, it is essential to recognize the substantial impact of agricultural management. This influence is not limited to the physical and chemical qualities of soil, but also extends to the composition and diversity of soil microbial communities (Schmidt et al., 2019). Previous studies have demonstrated that agricultural management significantly impacts the soil microbial community (Salles et al., 2006; Hartman et al., 2018b; Schmidt et al., 2019). A study on maize also showed that the influence of different nutrient additions on bulk soil can be extended into rhizosphere soil (Bakker et al., 2015). Our data corroborate these findings, showing that, in contrast to plant cultivar, the agricultural treatment applied in the soil profoundly impacts the composition of the rhizosphere microbial community (Fig. 2).

Although the above-mentioned findings demonstrate the necessity of considering the effects of treatments on soil microbial community structure in agricultural production, our data revealed that the impact of agricultural management was dependent on the microbial group targeted at the level of composition rather than diversity. In fact, although previous studies suggest that soil variables influence bacterial diversity more than fungal diversity (Wang et al., 2020). We observed no significant effect of treatment on microbial diversity (Figs. S3 and S4). Our results are in line with earlier findings that species composition was more susceptible to environmental factors than species richness (Morgan Ernest and Brown, 2001; Hartmann and Widmer, 2006). At the compositional level, we observed, for instance, that high-nutrient supply treatments (i.e., Fertiliser and Fertiliser plus Pesticide treatments) exhibited a more extensive disturbance in the bacterial community composition than other treatments (Fig. S9a). This observation aligns with previous research findings, demonstrating that fertilization can induce important alterations in bacterial diversity and composition of the maize rhizosphere (Wang et al., 2018). As expected, we observed that the fungal communities were more sensitive to the treatments involving pesticides (i.e., Pesticide and Fertiliser plus Pesticide treatments; Fig. S9b). These two treatments contain various fungicides that contribute to the observed variation in the fungal community. Collectively, our study highlights the differential responses of bacterial and fungal communities to agricultural treatments, emphasizing the importance of accounting for the soil microbiome communities when making decisions about agricultural practices and crop cultivation.

### 4.3. Biological management promotes co-occurrence networks and enhances the correlation between plants and the microbiome

The interactions between various soil microbial communities are intricate, including predation, competition, and mutualism (Faust and Raes, 2012). The interaction influences resource allocation, biodiversity, and ecological niches in soil systems. For instance, protists are consumers of bacteria and fungi, and any treatment effect on this microbial group potentially cascades into bacterial and fungal communities (Xiong et al., 2018). Bacteria and fungi also regulate each other, and form a web of cooperative and competitive relationships. They can collaborate to optimize nutrient uptake (Dellagi et al., 2020), and engage in antagonistic effects for substrate utilisation (Mille-Lindblom et al., 2006). In addition, fungal diversity can mediate the assembly of bacterial community in stressful environments (Jiao et al., 2022). Our data on networks revealed that biological practices promoted the co-occurrence of taxa and their interconnectivity (Fig. 4). In other words, our biological management selected microorganisms that responded in a similar way to the tested consortia, despite their differences. Although our methodology does not allow us to determine whether these interactions are indeed occurring in the soil, which requires non-destructive spatial sampling, it does indicate a higher probability. Considering that the stability of microbial networks increases with their connectivity and complexity (Yuan et al., 2021), our results could indicate that the higher microbial interactions promoted by biological management could lead to a soil microbiome less susceptible to external disturbances.

To further investigate the complex relationships between plant cultivar, agricultural management, the rhizosphere microbiome, and their influence on plant growth, we used piecewise structural equation modelling (SEM; Fig. 5), which has been widely used to explore the relationship between environmental factors, soil microbial community, plant development, and ecosystem function (Delgado-Baquerizo et al., 2016; Schmidt et al., 2019; Wu et al., 2022). These results highlighted the contradictory effects of biological and chemical management further. Regarding biological management, we showed that treatment had an effect on microbial composition for plant below-ground biomass. By compiling these results with those observed for the co-occurrence network data, we argue that biological management increases the interaction within microbial communities and promotes plant growth through the soil microbiome. Future studies focusing on the interaction between biological management practices and other crops selected for higher microbiome interactions are needed to confirm the generality of the additive effects observed in our study.

Conversely, our study showed that chemical management reduces the interaction within microbial communities and decreases the correlation between plant performance and the microbial community, removing the effect of the microbial community on plant growth. This confirms the previous research, emphasizing that under optimal conditions (i.e., high-nutrient or pathogen-free), the screening effect of the plant on microbiomes is declined, and the function of microbiomes on plant growth is disregarded (Wissuwa et al., 2009; Geisseler and Scow, 2014). Previous studies have also demonstrated that chemical management degrades the interaction between plants and microbes, which is harmful to the stability and balance of ecosystems (Huang et al., 2019; Molina-Santiago and Matilla, 2019). Overall, we argue that the plant cultivar and the rhizosphere microbiome collaborate to regulate plant growth under biological management. However, the rhizosphere microbial community decouples from plant growth under chemical management. These findings emphasize the benefits of plant-microbiome interaction on plant growth and highlight the essential role of management regimes in shaping these relationships.

Our study faced an unexpected challenge in the form of *Phytophthora infestans* occurrence in the field, which disrupted our analyses after the seventh week. Consequently, the results presented in this study are confined to the early stages of plant development (before flowering). The selected cultivars were insufficient to control *Phytophthora*, although some plants did exhibit resistance. We suggest that the identification of genetic traits associated with the cultivar Salto could be implemented in future breeding programs where *Phytophthora* resistance is also considered. Due to the complexity of the field, we did not collect plant root exudates in this study. The absence of this data component does statistically reduce the strength of the correlation between plant cultivars and the composition of rhizosphere microbial communities. It has been proved that the maize genotype profoundly influences the rhizosphere microbial community in long-term experiments (Aira et al., 2010; Walters et al., 2018). This finding highlights the importance of considering the specific context and crop stage when assessing the effects of treatments on soil microbial communities. To address this gap, future studies should extend the experiment duration and examine root exudates to explore how plant cultivars with different functional traits affect the microbial communities through their metabolites.

## Conclusion

Our findings reveal that plant cultivars possessing high MITs exhibit superior plant growth performance when compared to the commercial cultivar Desiree. This enhanced performance is potentially attributed to the improved interactions between plants and the rhizosphere microbiome. Agricultural management serves as the primary factor influencing the composition of the rhizosphere microbial community. More specifically, biological management increases inter-kingdom interactions. In this context, both potato cultivar and the rhizosphere microbiome collaborate synergistically to regulate and enhance plant growth. Conversely, chemical management practices lead to a decoupling of the rhizosphere microbial community from plant growth. In summary, our study highlights the pivotal role of plant cultivar, agricultural management, and microbiome interactions in shaping plant development. It emphasizes the potential benefits of plant cultivars with MITs and the positive impacts of biological management.

## Availability of Data

The raw sequencing data are available in the National Center for Biotechnology Information (NCBI) Sequence Read Archive (SRA) under the accession number PRJNA1041735.

## Author contributions

J.F.S. and J.T.M.E. designed the experiment; T.Z. and X.J. performed the experiment, laboratory work and data analysis; X.L. participated in field work and data analysis; J.N., E.A., and R.G. took care of the sequencing of soil DNA extraction samples; K.T., A.P. and D.M. provided all the potato tubers; G.B., F.S., and T.C. developed the consortium for biological management; T.Z. wrote the manuscript. All the authors contributed to the modification and approval of the final version.

## Acknowledgements

This study was funded by ERA-NET Cofund SusCrop project potatoMETAbiome, which is supported by the European Union’s Horizon 2020 research and innovation program (grant agreement No 771134; French National Agency, grant number -ANR-18-SUSC-0001) and part of the Joint Programming Initiative on Agriculture, Food Security and Climate Change (FACCE-JPI). We would like to acknowledge Averis Seeds BV’s support in conducting the field trial. Special thanks to Johan Hopman, Hendrik Jan Schepel, Nico Rookmaker and Natasja Dammer. We appreciate Pier Oosterkamp, Sjoukje Hoekstra and Zhilei Gao from ECOstyle for providing the microbial strains used in the Consortium BP treatment. We thank Jolanda K. Brons, Michael Cen Feng, Jan Veldsink, Stefanie Vink, Pim Henstra, Siyu Mei, Pina Brinker, Panji Cahya Mawarda, Edisa Garcia Hernandez, Qian Chen, Yanfang Wang and Fanny Guedea for their help in the fieldwork as well as Maria Fedczak and Alicja Przewłoka in seed tuber production. T.Z. and X.L. were supported by a scholarship from the China Scholarship Council (CSC) and the University of Groningen scholarship program. J.N. was supported by a scholarship from the Local Council of Pyrénées-Atlantiques, E2S UPPA and the University of Groningen scholarship program. We thank PGTB for sequencing (Genome Transcriptome Platform of Bordeaux, doi:10.15454/1.5572396583599417E12) (Univ. Bordeaux, INRAE, BIOGECO, F-33610 Cestas, France) and notably Erwan Guichoux and Emilie Chancerel.

