## Supplement for "Biological management, rather than chemical management, promotes the interaction between plants and their microbiome"

### Supplementary information

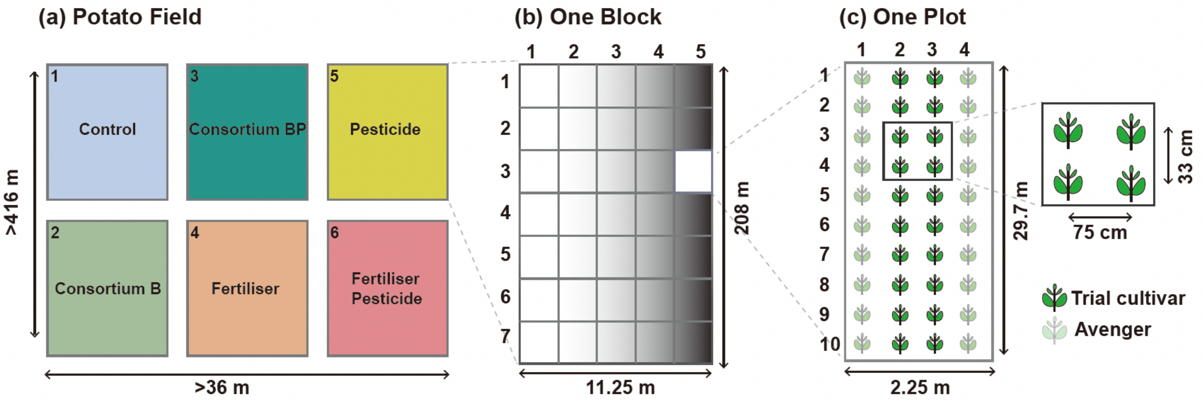


**Fig. S1 The Schematic plot represents the field trial setup.** (A) The distribution of six treatments in the field trial. (B) In each treatment/block, three replicates/plots of eleven potato cultivars are distributed in 35 plots (5 × 7). Potato cultivar Avenger fills up blocks in which the plots were not distributed to the trial cultivars. (C) In each plot, 40 plant individuals (10 × 4) are planted, in which 20 individuals in the middle two columns will be the trial cultivars. Plants in the two outside columns are Avenger, which are used to reduce edge effects. Within a plot, potatoes are planted at 75 cm row spacing and 33 cm within-row spacing. In the end, four cultivars are removed due to insufficient tubers, seven cultivars were kept for analysis.

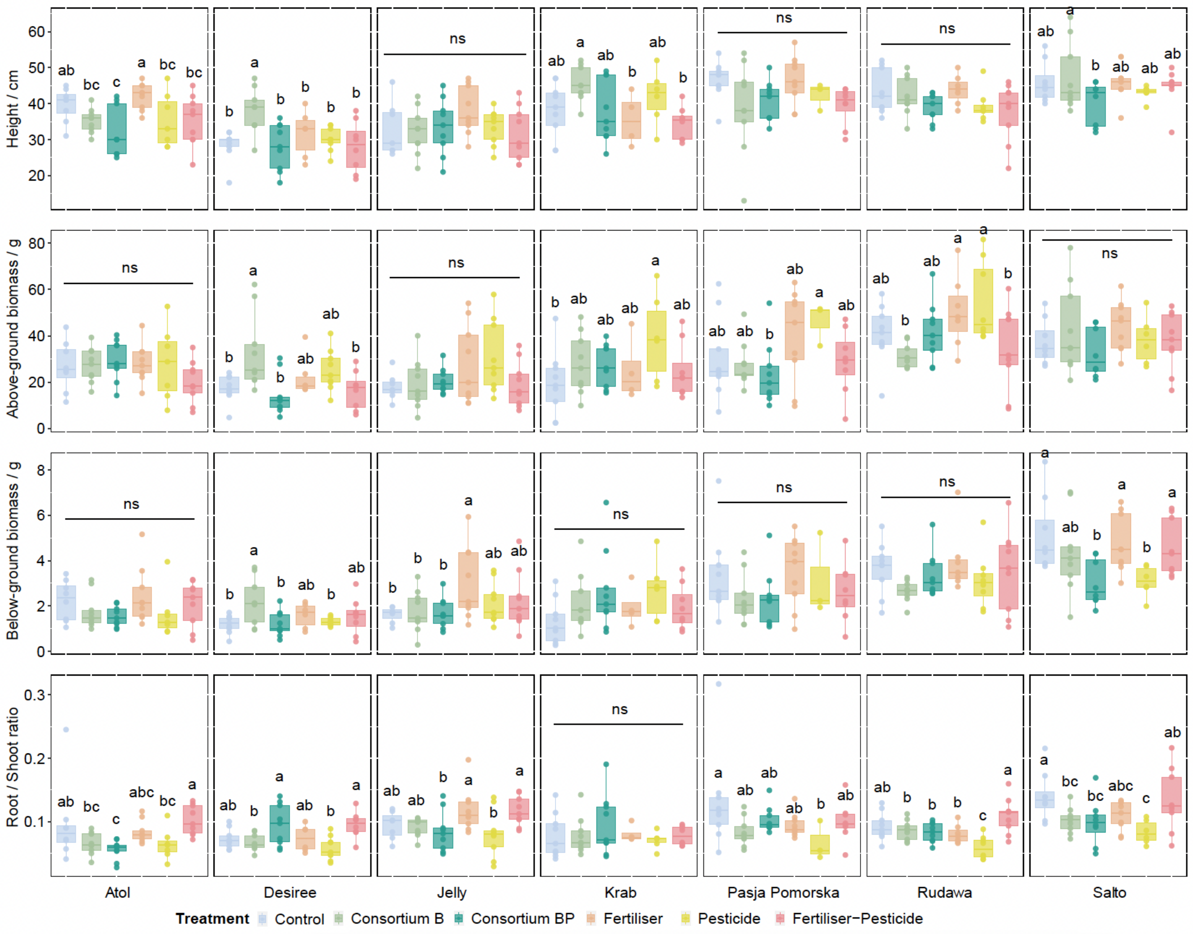


**Fig. S2 Plant performance of six treatments across the same cultivars.** Letters indicate significant differences across treatments (Duncan post hoc test).

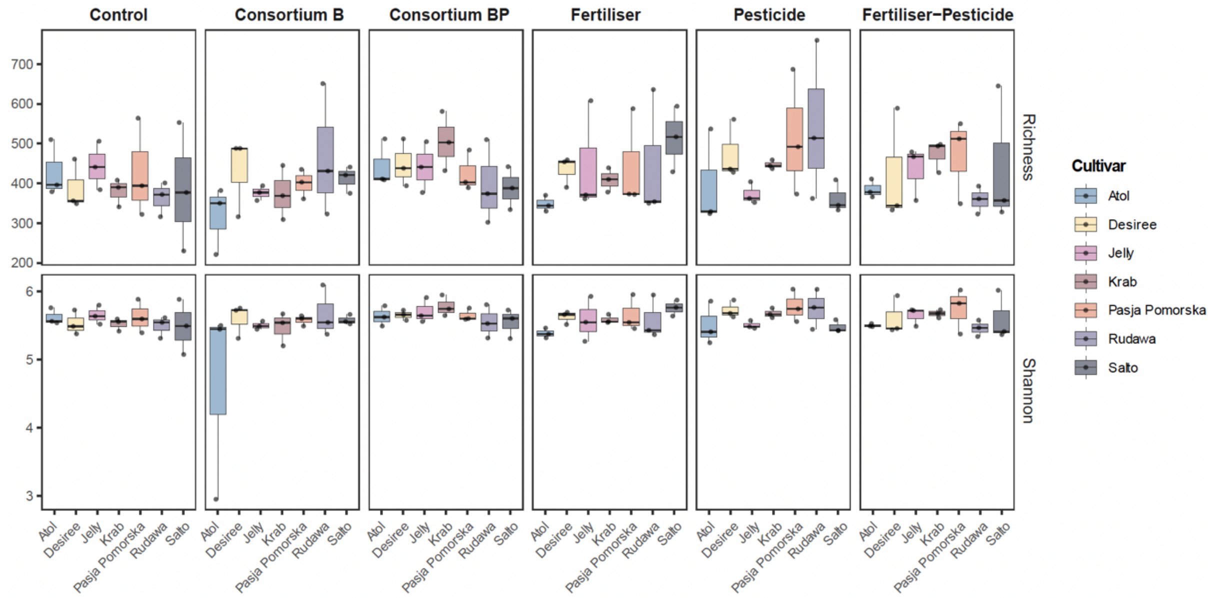


**Fig. S3 Alpha diversity of bacterial community of seven cultivars across treatments.** No significant differences across cultivars (Tukey post hoc test).

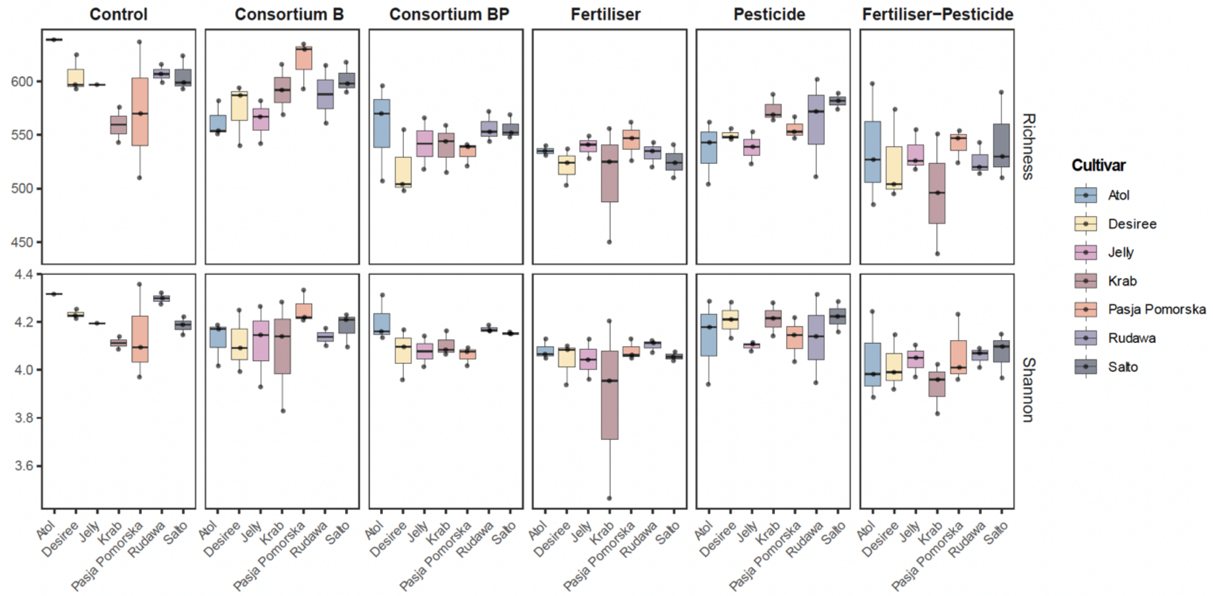


**Fig. S4 Alpha diversity of fungal community of seven cultivars across treatments.** No significant differences across cultivars (Tukey post hoc test).

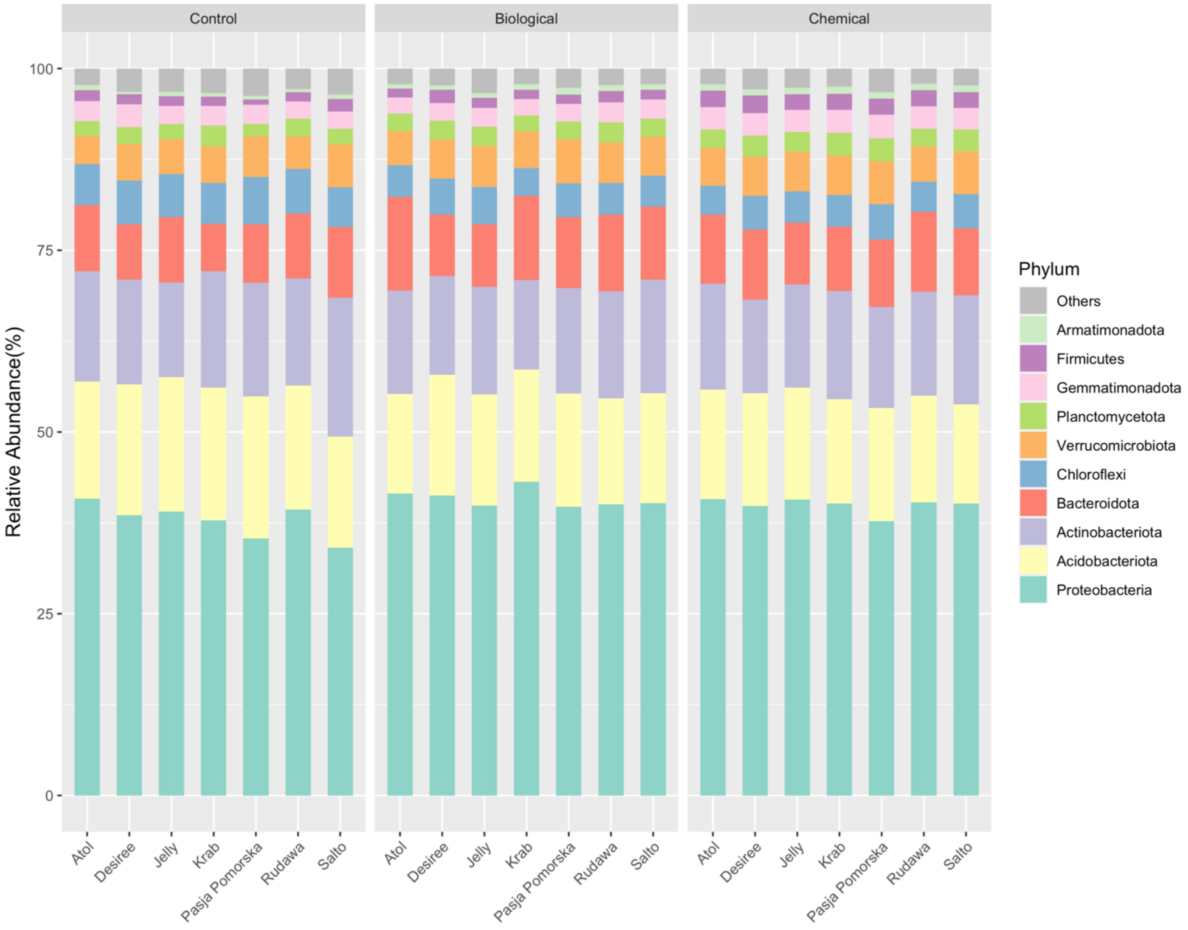


**Fig. S5 Taxonomy classification of bacterial ASVs at phylum levels.** The top 10 most relative abundance are shown in the figure. Other phyla are assigned as "Others".

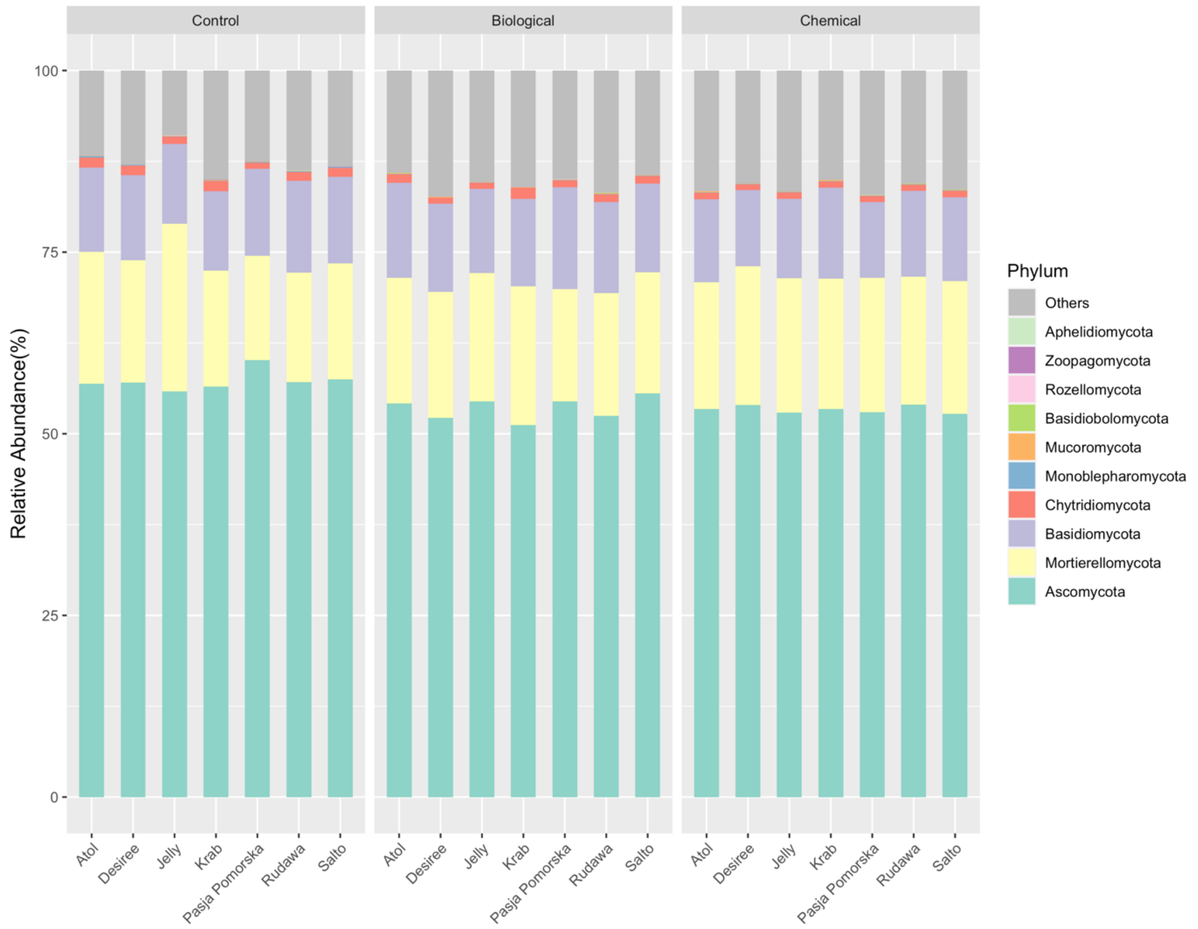


**Fig. S6 Taxonomy classification of fungal OTUs at phylum levels.** The top 10 most relative abundance are shown in the figure. Other phyla are assigned as "Others".

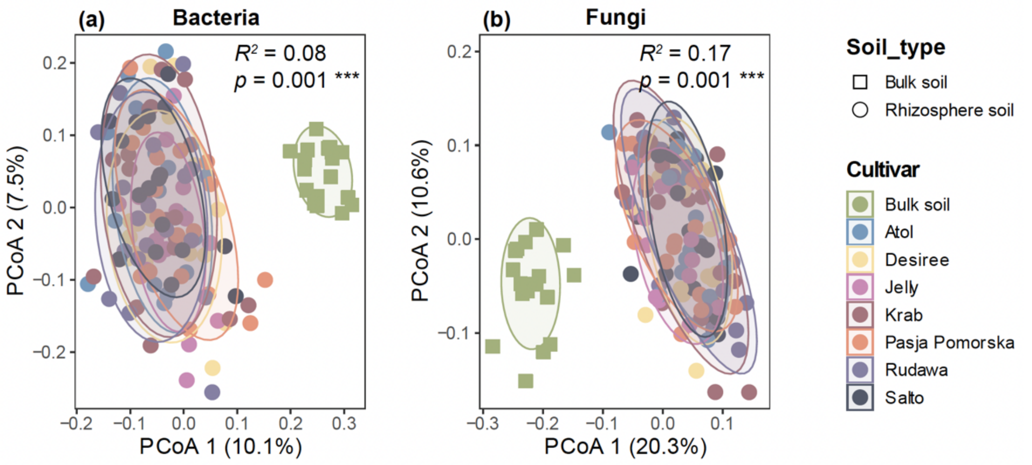

**Fig. S7 The distribution of bulk and rhizosphere soil microbial communities, displayed separately for bacteria and fungi, were visualised via a principal coordinates analysis (PCoA) based on Bray-Curtis matrices.** Colours indicate cultivars and bulk soil. Shapes indicate soil type. The PERMANOVA (Adonis) results show the influence of soil type on community composition. $R^2$ explains variation; $p = 0.001$ indicates the significant difference in microbial community composition in different soil compartments.

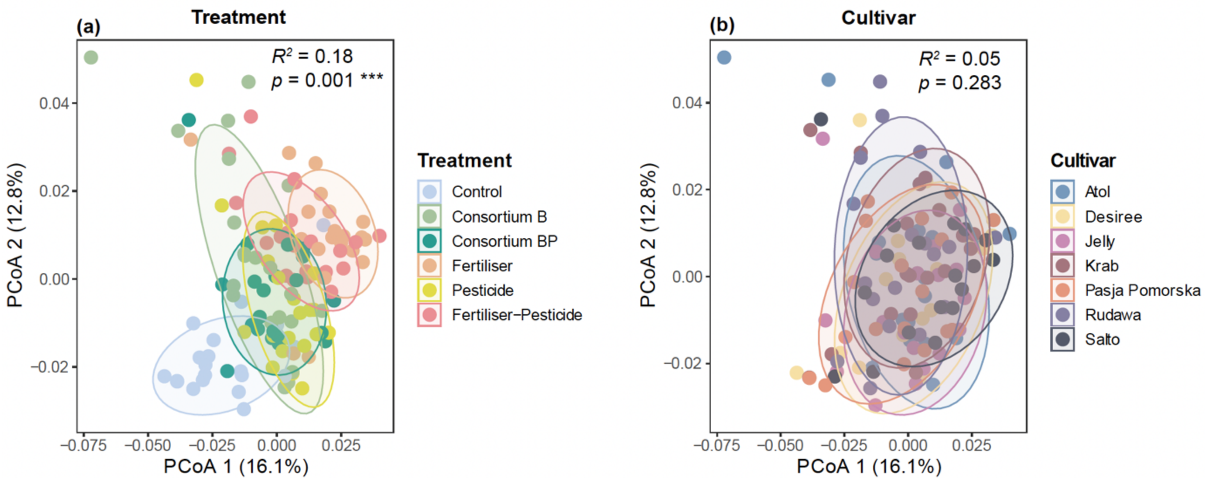


**Fig. S8 The effect of treatment and cultivar on the composition of rhizosphere bacterial community, were visualised via a principal coordinates analysis (PCoA) based on weighted UniFrac dissimilarity matrices.** Colours in the right panel indicate treatments. Colours in the left panel indicate cultivars. PERMANOVA (Adonis) results show the influence of factors on community composition. $R^2$ explains variation, and $p < 0.05$ means the significant difference in microbial community composition in treatments (right panel) and cultivars (left panel).

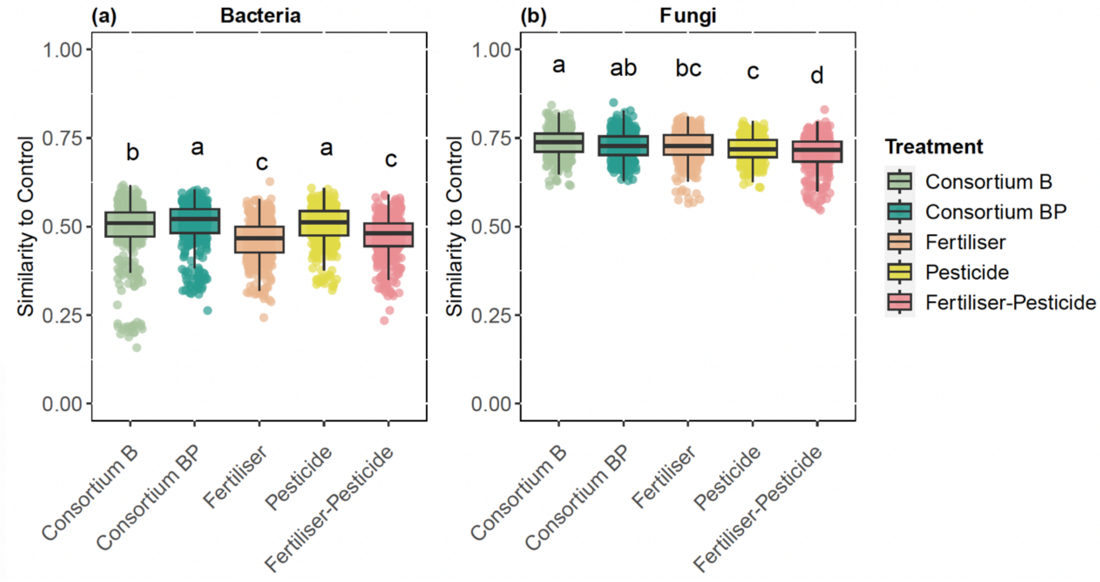


**Fig. S9 The similarity of treatment on the distance of rhizosphere soil microbial communities, compared to Control.** Bacteria and fungi were visualised separately based on similarity, which is 1−Bray-Curtis matrices. Letters indicate significant differences across treatments (Tukey's Test).

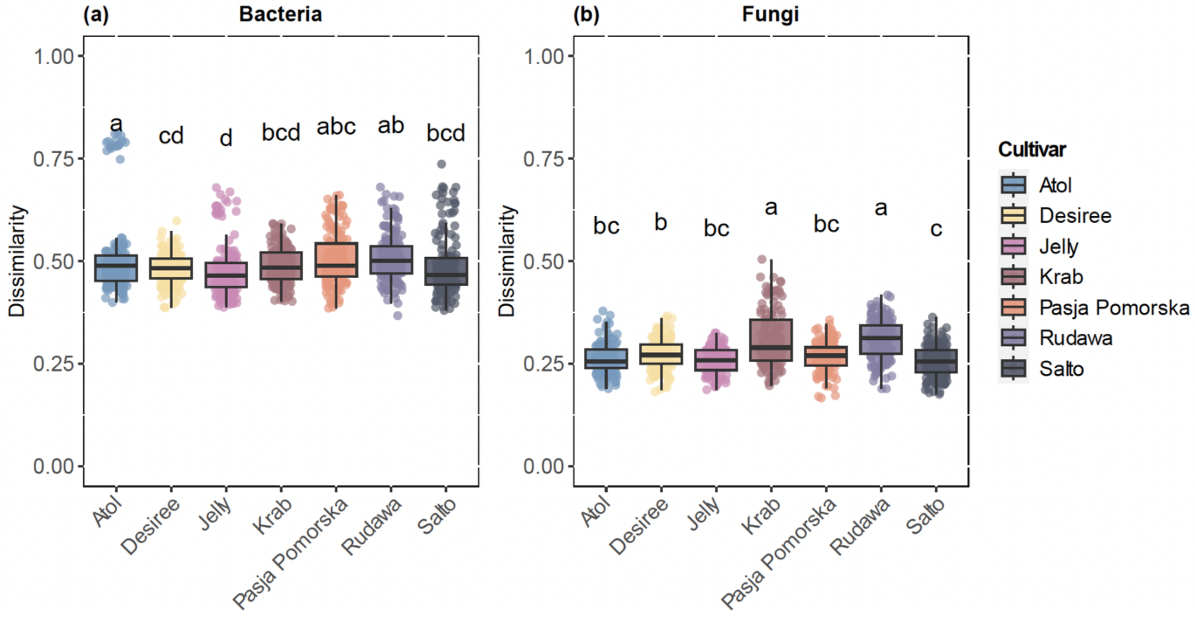

**Fig. S10 The dissimilarity distance of rhizosphere soil microbial communities within cultivars.** Displayed for bacteria and fungi separately were visualised based on Bray-Curtis matrices. Letters indicate significant differences across cultivars (Tukey's Test).

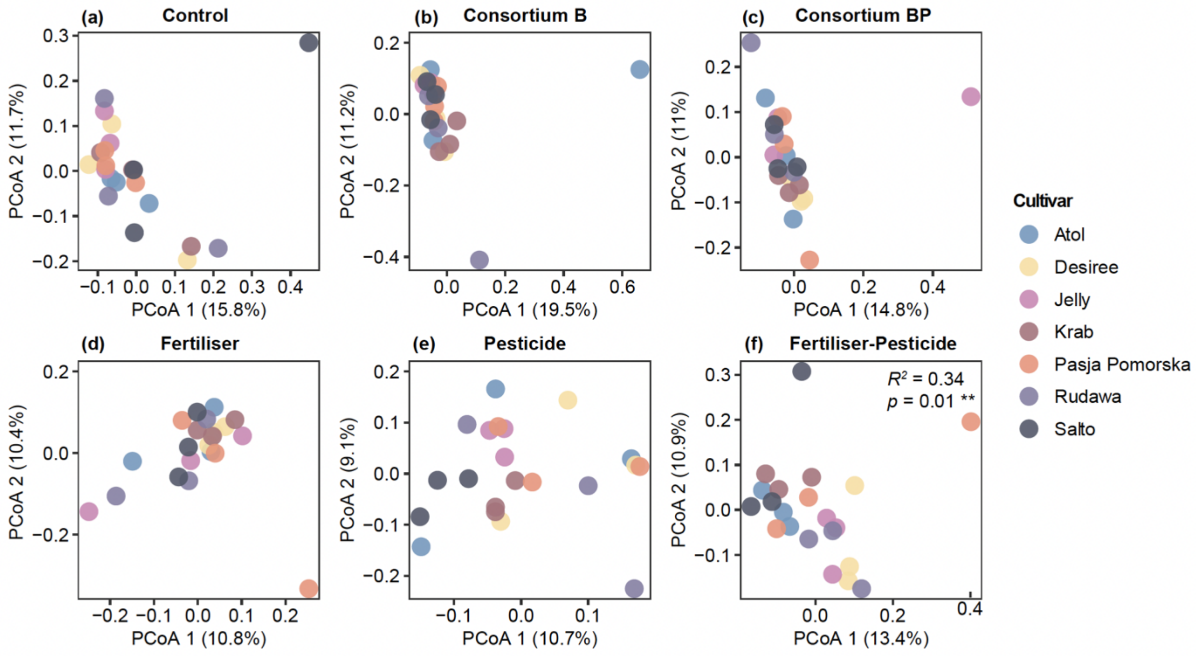


**Fig. S11 The effect of cultivar on the composition of rhizosphere bacterial microbial community, displayed for different treatments, were visualised via a principal coordinates analysis (PCoA) based on Bray-Curtis matrices.** Colours indicate cultivars. PERMANOVA (Adonis) results show the influence of cultivar on bacterial community composition. $R^2$ explains variation, and $p < 0.05$ means the significant difference in bacterial community composition in treatments. The panels without a PERMANOVA (Adonis) result means that the cultivar does not significantly affect bacterial community composition.

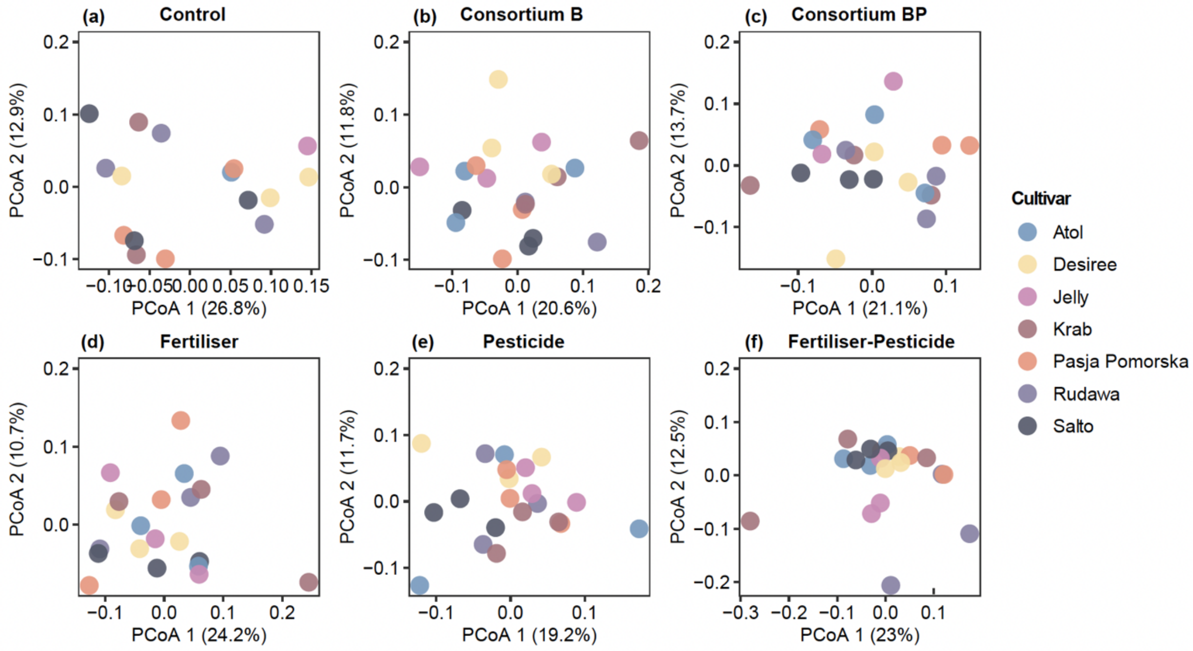


**Fig. S12 The effect of cultivar on the composition of rhizosphere fungal microbial community, displayed for different treatments, were visualised via a principal coordinates analysis (PCoA) based on Bray-Curtis matrices.** Colours indicate cultivars. The panels without a PERMANOVA (Adonis) result means that the cultivar does not significantly affect fungal community composition.

**Table S1. Chemical compounds in the Pesticide treatment**

| Commercial name | Active component | Concentration |
|---|---|---|
| Amphore flex | cymoxanil | 18.00 % |
| | mandipropamid | 25.00 % |
| Arcade | metribuzin | 80.00 g/l |
| | prosulfocarb | 800.00 g/l |
| Basagran | bentazon | 480.00 g/l |
| Carial star | difenoconazool | 250.00 g/l |
| | mandipropamid | 250.00 g/l |
| Curzate partner | cymoxanil | 60.00 % |
| Gazelle | acetamiprid | 20.00 % |
| Infinito | fluopicolide | 62.50 g/l |
| | propamocarb | 625.00 g/l |
| Narita | difenoconazool | 250.00 g/l |
| Ranman top | cyazofamide | 160.00 g/l |
| Sencor sc | metribuzin | 600.00 g/l |
| Signum | boscalid | 26.70 % |
| | pyraclostrobine | 6.70 % |
| Titus | rimsulfuron | 25.00 % |
| Zorvec endavia | benthiavalicarb-isopropyl | 70.00 g/l |
| | oxathiapiproline | 30.00 g/l |
